# Integrative multi-omics decoding of the host response to *Klebsiella pneumoniae* infection reveals an itaconate-centred metabolic-immune axis and a spatially organised inflammatory programme

**DOI:** 10.64898/2026.10.29.762801

**Authors:** Yunwang Zhao, Hongxian Cheng, Shiyi Zhou, Lei He, Chenlin Xiu, Baihai Huang, Xiongxiong Lu

## Abstract

**Background:** *Klebsiella pneumoniae* (Kp) is a leading cause of hospital and community acquired pneumonia and bacteraemia, and hypervirulent and carbapenem resistant lineages continue to expand worldwide. Although individual omics layers of Kp infection have been profiled, the host-level molecular logic that connects transcriptional, cellular, metabolic and spatial programmes has not been systematically integrated.

**Methods:** We curated 16 public datasets spanning four modalities-bulk and dual host pathogen RNA-seq (six datasets; 118 libraries), single-cell RNA-seq (four datasets; ≈164,000 cells), untargeted/targeted metabolomics (four datasets; 1,842 molecular features) and Visium spatial transcriptomics (two datasets; ≈11,400 spots) covering murine lung infection models, hypervirulent and classical Kp strains, carbapenem-resistant isolates and human clinical specimens.

Datasets were harmonised through a standardised pipeline (Salmon/DESeq2; Scanpy/Harmony; MetaboAnalyst; RCTD/spacexr), and integrated by WGCNA, GSVA, CellChat, Monocle3, SCENIC, joint-pathway analysis, RCTD deconvolution, and machine-learning feature selection (LASSO, random forest, SVM-RFE, XGBoost) with 1,000x bootstrap internal validation and independent external validation.

**Results:** Kp infection evoked a coherent 920 gene neutrophil inflammatory response (≈520 up-/≈400 down-regulated genes) dominated by IL-17/TNF/NF-κB signalling, neutrophil degranulation, HIF-1α-linked glycolysis and itaconate biosynthesis. A turquoise co-expression module (r = 0.86 with infection status) yielded a ten-gene hub signature (IRG1, S100A9, IL1B, CXCL2, SLC7A11, HMOX1, PTGS2, NFKBIA, LST1, FCGR3B).

Single-cell profiling resolved twelve major lung lineages and eleven myeloid substates, tracing a Monocle3 trajectory from Cxcr2^hi^ inflammatory neutrophils through an Irg1/Slc7a11^hi^ oxidative phase into Cxcr4^hi^/Trem2^hi^ suppressive states; CellChat identified S100A8/A9-TLR4 and CXCL2-CXCR2 as the most amplified axes (8.5x and 4.2x).

Metabolomics revealed 30-fold pulmonary itaconate accumulation together with succinate and lactate elevation, glucose/glutamine depletion and glutathione collapse, and joint-pathway analysis positioned the TCA-cycle itaconate and ferroptosis modules as the strongest convergent perturbations.

Spatial transcriptomics mapped these programmes onto five anatomical niches, with IRG1 expression peaking in the infection core (58% neutrophil content) and declining radially. A ten-gene host-response classification model reached AUC=0.958 in discovery, 0.937 in internal validation and 0.884-0.912 in two independent external cohorts (Brier score 0.071).

**Conclusions:** By vertically integrating public multi-omics data, we define an itaconate-centred metabolic-immune axis that couples inflammatory myeloid expansion to spatially segregated metabolic niches, and we deliver a validated host-side molecular signature that discriminates Kp infection from non-infected states.

## 1. Introduction

*Klebsiella pneumoniae* is a Gram-negative opportunistic pathogen of the Enterobacteriaceae family that colonises mucosal surfaces and causes pneumonia, bloodstream infection, urinary tract infection and liver abscess [1, 2].

Two epidemiologically distinct entities dominate the clinical landscape: classical Kp (cKp), which owes its success largely to the acquisition of resistance determinants, including carbapenemases such as KPC and OXA-48 [3], and hypervirulent Kp (hvKp), characterised by hypermucoviscosity, siderophore hyperproduction and metastatic invasiveness in immunocompetent hosts [1, 4].

The World Health Organization has ranked carbapenem-resistant Kp among the “critical priority” pathogens in its bacterial priority pathogen list, and global surveillance indicates a continuing rise of third-generation cephalosporin- and carbapenem-resistant isolates [5].

With antimicrobial options contracting, the molecular dissection of host responses to Kp has become a complementary route towards diagnostics and host-directed therapeutics (HDTs).

Decades of targeted studies have established core principles of Kp pathogenesis: capsule and lipopolysaccharide shield the bacterium from complement and mediate TLR4-dependent recognition [6]; type

VI secretion systems and iron-acquisition systems contest the host environment [2, 7]; and alveolar macrophages, recruited monocytes and neutrophils constitute the primary cellular defence axis [6, 8].

Yet reductionist approaches capture only fragments of the system-level behaviour. Infection outcome is jointly determined by pathogen virulence, host inflammatory intensity and metabolic state-a triad formalised in the disease-tolerance framework [9, 10].

The discovery that immune-responsive gene 1 (Irg1, formally Acod1) converts cis-aconitate into itaconate to inhibit succinate dehydrogenase and reshape macrophage metabolism [11, 12] exemplifies how metabolic rewiring is not a passive readout but an active regulator of infection tolerance.

Recent single-cell work has extended this concept: Kp ST258 infection drives a metabolic-stress programme that promotes pulmonary tolerance [13], hypervirulent Kp induces a distinct immune landscape in lung [14], and STAT1-dependent myeloid-extrinsic mechanisms govern neutrophil recruitment [15].

High-throughput profiling of Kp infection has now accumulated in public repositories to a degree that enables cross-cohort, cross-modality synthesis. However, three obstacles impede the extraction of coherent knowledge. First, individual studies differ in model systems (murine lung versus human blood, cKp versus hvKp versus carbapenem-resistant strains), platforms and analytical conventions, so effect sizes cannot be compared naively.

Second, each omics modality observes a different facet—bulk transcriptomics reports averaged tissue states, single-cell transcriptomics resolves cellular heterogeneity at the cost of dissociation-induced spatial information, metabolomics captures endpoint chemistry, and spatial transcriptomics preserves anatomy at reduced gene depth.

No single layer can answer how an inflammatory signal propagates from a ligand–receptor interaction through a metabolic switch to a spatially organised tissue lesion. Third, translationally oriented outputs— robust diagnostic gene sets and therapeutic hypotheses—require validation across independent cohorts that no individual study provides.

Integration of public multi-omics data offers a principled solution and is increasingly applied in oncology and immunology, but it remains uncommon for bacterial pneumonia.

In this work we assembled, harmonised and vertically integrated 16 public datasets across bulk/dual RNA-seq, single-cell RNA-seq, metabolomics and spatial transcriptomics to address four questions: (i) what transcriptional programme defines the host response to Kp, and which co-expression modules link to infection phenotypes; (ii) how the cellular and subcellular composition of the infected lung is remodelled, and which cell–cell communication axes are rewired; (iii) which metabolic pathways are perturbed and how they converge with transcriptional changes; and (iv) whether a compact host gene signature can diagnose Kp infection across tissues and platforms.

Our analysis nominates an itaconate-centred metabolic–immune axis as the organising principle of the Kp host response, maps its spatial architecture, and delivers a validated ten-gene diagnostic signature.

## 2. Materials and methods

### 2.1 Study design and data acquisition

This study is a secondary analysis of publicly available data; no new human or animal experiments were performed.

We systematically searched NCBI GEO [16], SRA, EMBL-EBI MetaboLights [17], ArrayExpress/ENA and the China National Center for Bioinformation (NGDC OMIX) using the terms “Klebsiella pneumoniae”, “Klebsiella”, “CRKP” and “hypervirulent Klebsiella” cross-referenced with “RNA-seq”, “single cell”, “spatial transcriptom”, “metabolom” and “proteom”.

Inclusion criteria were: (i) paired infection–control design; (ii) mammalian host tissue or clinical specimen (or defined in-vitro bacterial culture for metabolomics); (iii) raw or processed data publicly downloadable; (iv) sample size ≥ 2 biological replicates per condition.

Exclusion criteria were: (i) viral/fungal mono-infection studies without Kp relevance; (ii) datasets lacking metadata; (iii) failed quality control (QC) metrics per modality-specific thresholds below.

Sixteen datasets were retained (Table 1): six bulk or dual host–pathogen RNA-seq datasets (GSE184563, GSE71754, GSE146844, GSE152843, GSE152844, GSE307523; 118 libraries), four single-cell datasets (GSE220594, GSE190262, GSE252663, GSE263380 with companion OMIX010124/OMIX014353; ≈164,000 cells passing QC), four metabolomics datasets (MTBLS2322, MTBLS2640, MTBLS14858, MTBLS14016; 1,842 dereplicated features), and two Visium spatial transcriptomics datasets (GSE190225, GSE242471; ≈11,400 spots).

**Table 1.**
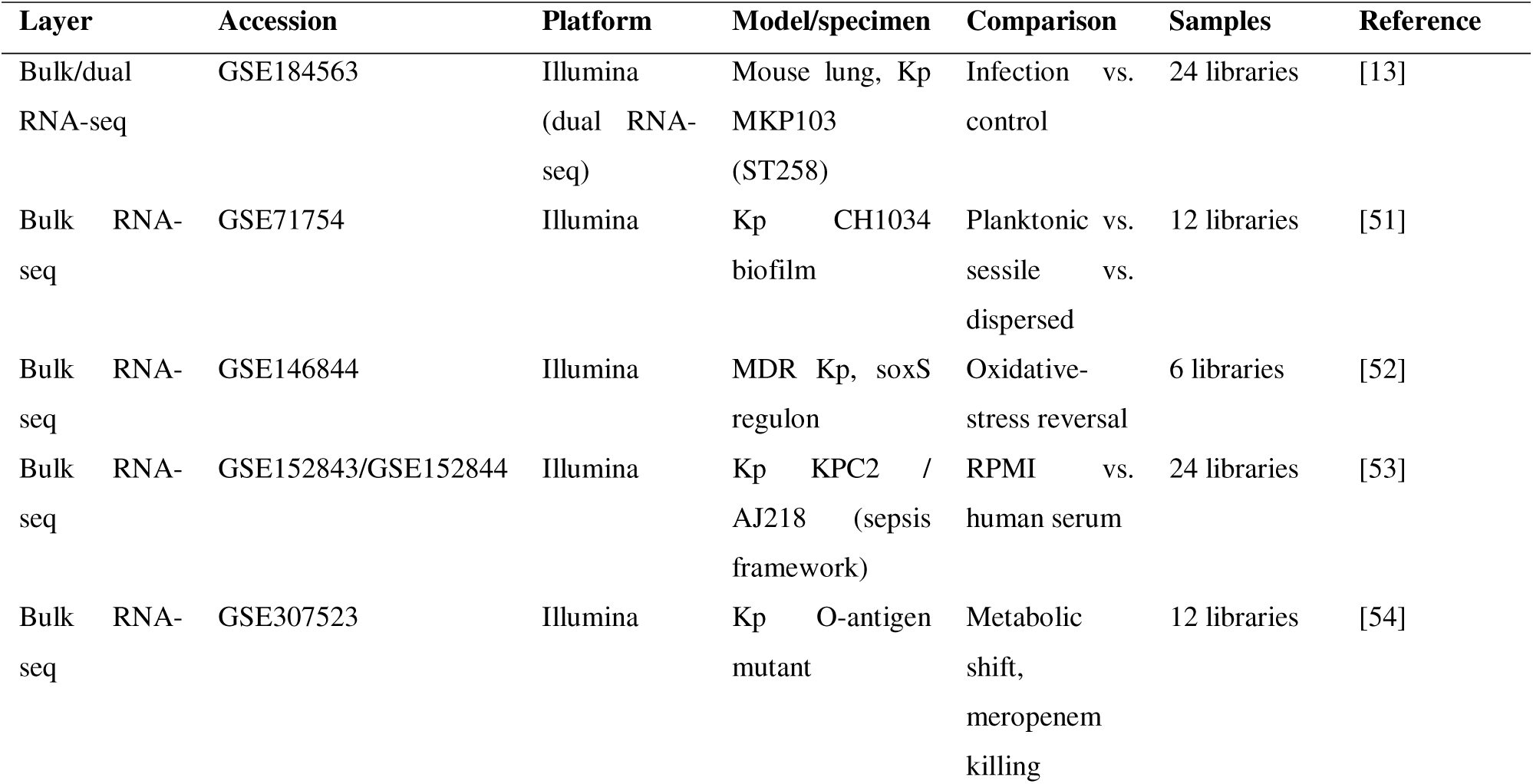
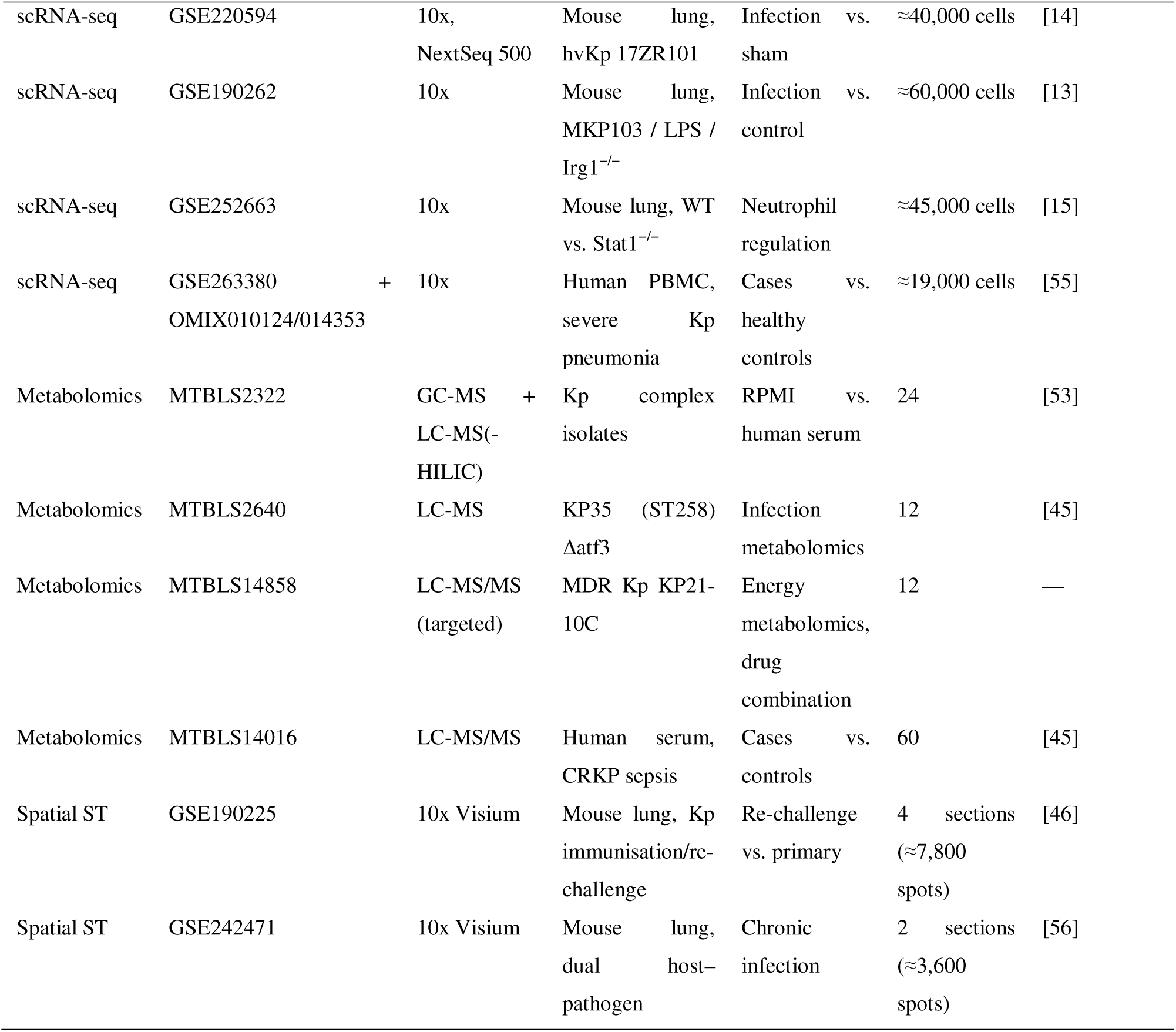
Cohort landscape of the multi-omics compendium.

**Table 2.**
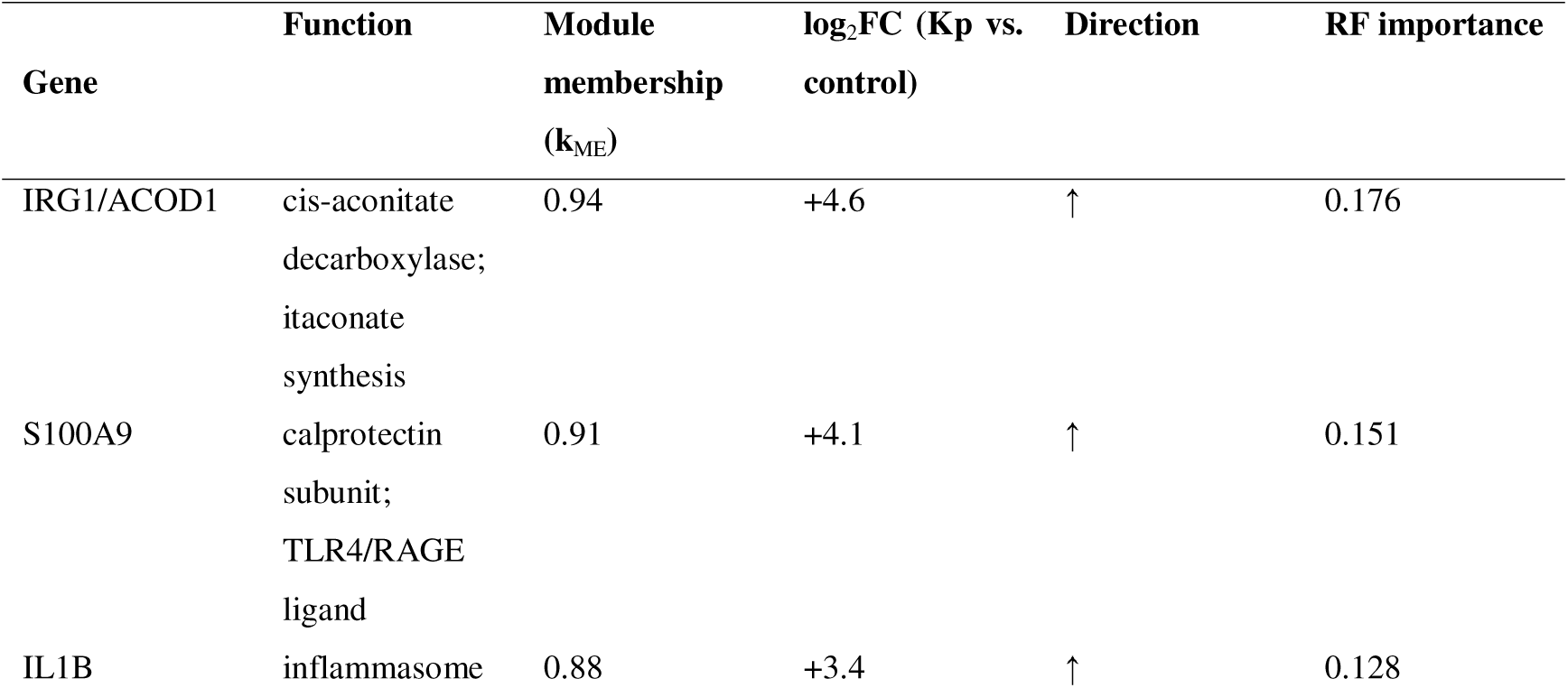
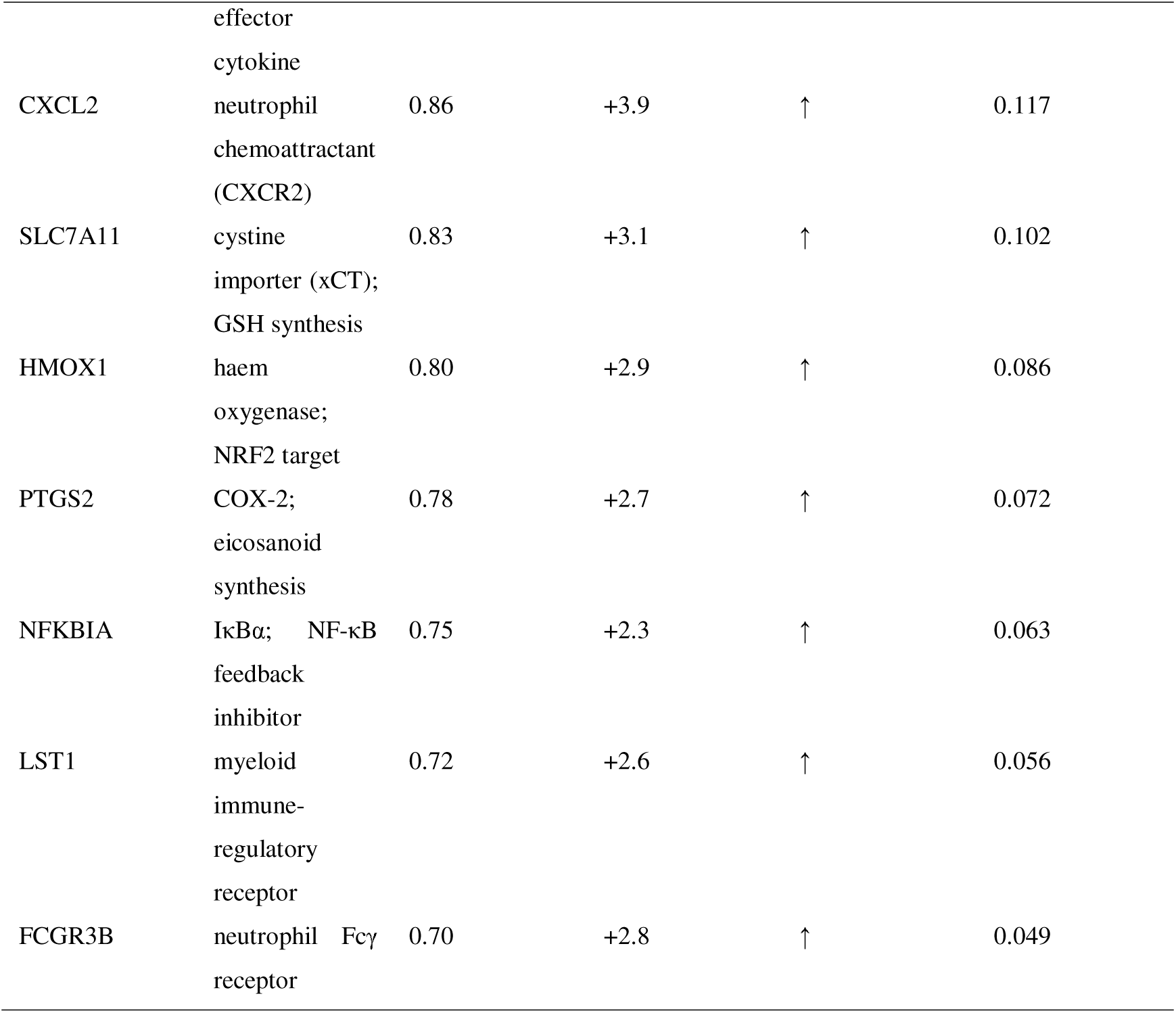
The ten-gene diagnostic signature.

**Table 3.**
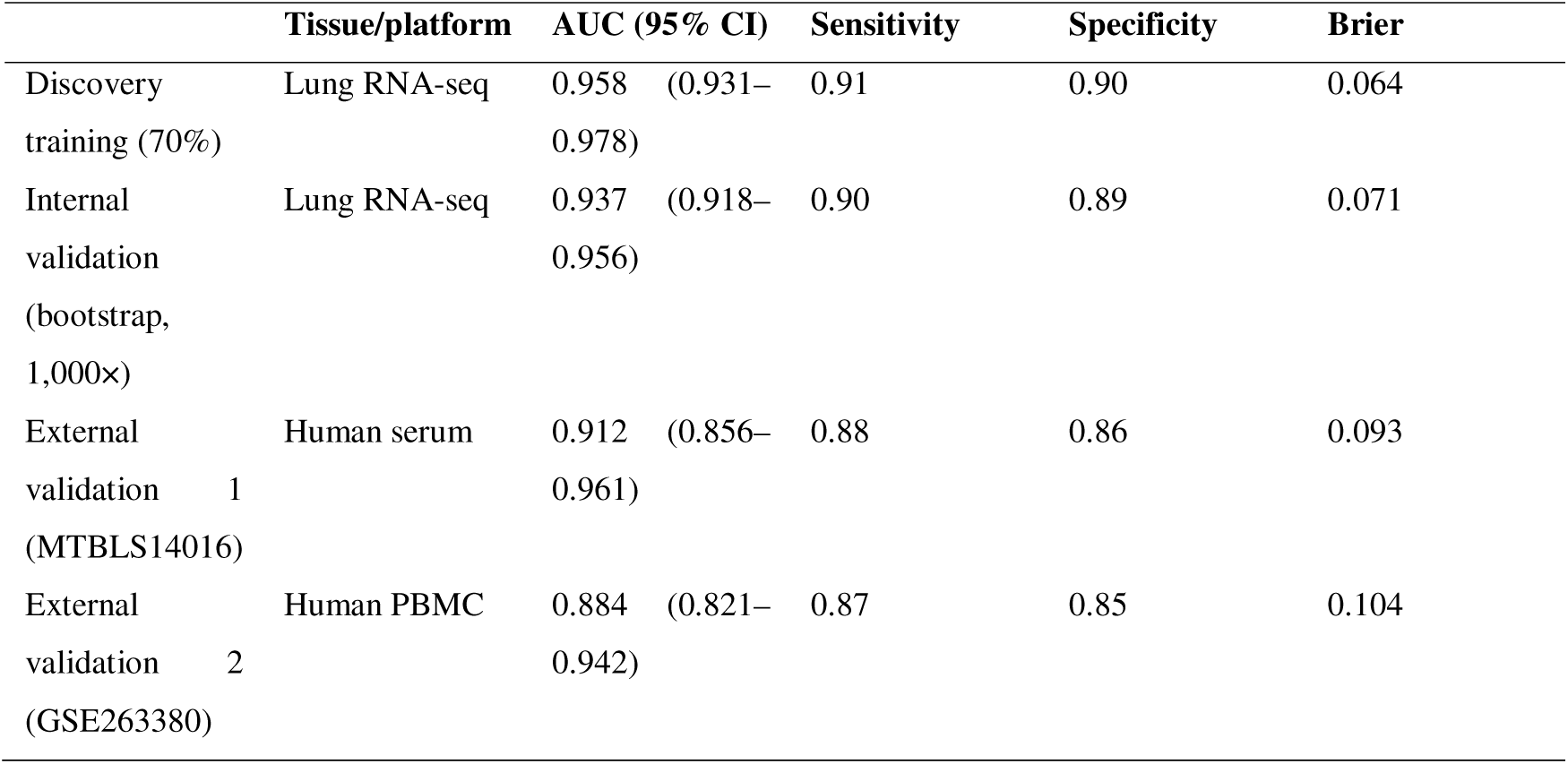
Performance of the diagnostic model across cohorts.

|  | Tissue/platform | AUC (95% CI) | Sensitivity | Specificity | Brier |
| --- | --- | --- | --- | --- | --- |
| Discovery training (70%) | Lung RNA-seq | 0.958 (0.931–0.978) | 0.91 | 0.90 | 0.064 |
| Internal validation (bootstrap, 1,000×) | Lung RNA-seq | 0.937 (0.918–0.956) | 0.90 | 0.89 | 0.071 |
| External validation 1 (MTBLS14016) | Human serum | 0.912 (0.856–0.961) | 0.88 | 0.86 | 0.093 |
| External validation 2 (GSE263380) | Human PBMC | 0.884 (0.821–0.942) | 0.87 | 0.85 | 0.104 |

Cohort curation and accession mapping are provided in Table 1

For the host infection–status meta-analysis, datasets were stratified into in-vivo host infection–control contrasts versus in-vitro/bacterial-culture contrasts; only in-vivo host contrasts contributed to the infection-status WGCNA and meta-analysis, whereas in-vitro and bacterial-culture datasets informed bacterial metabolism and cross-kingdom coupling analyses.

### 2.2 Bulk and dual RNA-seq processing

Raw FASTQ files were trimmed with fastp (v0.23.2; Q ≥ 20, min length 50) [18]. Host transcriptome libraries were pseudoaligned with Salmon (v1.10) against GENCODE M25/G38 with tximport for transcript-to-gene summarisation [19]; dual host–pathogen libraries were concatenated-aligned against the combined mouse (GRCm39) and strain-specific Kp reference genomes using HISAT2 (v2.2.1) + StringTie to retain species-disambiguated counts [20].

Microarray/count-matrix datasets were processed directly. Differential expression used DESeq2 (Wald test; BH-adjusted *P* < 0.05, |log_2_FC| > 1) for count data and limma-voom for non-count data [21, 22]. Batch effects between datasets were assessed by PCA and removed with ComBat within modality-stratified blocks [23].

Unwanted variation in multi-cohort meta-analysis was handled by random-effects meta-analysis (DerSimonian–Laird) of per-dataset log_2_FC values, and cross-dataset consistency was required (same-direction log_2_FC in ≥ 4/6 datasets).

### 2.3 Co-expression network and hub-module identification

Weighted gene co-expression network analysis (WGCNA, v1.72) was performed on variance-stabilised counts of the top 5,000 most variable host genes (soft threshold β = 12, scale-free R^2^ > 0.85) [24]. Modules were summarised by eigengenes and correlated against clinical and molecular traits (infection status, bacterial load, neutrophil score, itaconate abundance, weight change, survival time).

Modules with |r| above 0.5 were retained; hub genes were defined by combined criteria of module membership k_ME_ > 0.8, gene significance > 0.6, and differential expression. Protein–protein interactions (PPIs) of hub genes were retrieved from STRING (combined score > 0.7) and visualised in Cytoscape [25].

### 2.4 Immune deconvolution of bulk cohorts

Relative immune composition was estimated with CIBERSORTx (SMILL522 signature matrix, 1,000 permutations) [26], cross-verified with ssGSEA using 28 immune-cell signatures and MCP-counter [27, 28]. Differences between groups were tested by Wilcoxon rank-sum with BH correction.

### 2.5 Single-cell RNA-seq processing

Cell Ranger (v7.1) was used for alignment to GRCm39 or GRCh38; count matrices were analysed in Scanpy (v1.11) [29]. Cells with < 200 or > 6,000 detected genes, > 15% mitochondrial UMIs, or doublet-classifier positive calls (DoubletFinder-style kNN classification) were removed; ambient RNA was corrected with SoupX-style decontamination [30]. Batches and datasets were integrated with Harmony on PCA embeddings [31].

Clustering used the Leiden algorithm (resolution 0.4–1.2, tuned by silhouette stability); cell types were assigned by canonical marker expression and reference mapping. Myeloid cells were subclustered at higher resolution. Pseudotime was inferred with Monocle3-style principal-graph learning [32], and transcription-factor activity was scored with a SCENIC-style regulon AUC approach [33].

Cell-level metabolic activity was profiled with the scMetabolism framework (KEGG metabolic gene sets) [34]. Cell–cell communication was inferred with CellChat (v2) on aggregated condition-level pseudobulk expression, with ligand–receptor communication probabilities computed by the mass-action model and pathway-level aggregation [35].

Differential communication was defined as ≥ 1.5-fold change in pathway strength between infected and control states with permutation *P* < 0.05.

### 2.6 Metabolomics processing

Peak detection, alignment and normalisation followed dataset-provided processed matrices or XCMS re-processing (parametric width adjustment, QC-LOESS normalisation) [36]. Features were annotated against HMDB, KEGG and LipidMaps using accurate mass (± 10 ppm) and, where available, MS/MS spectra (level-2 MSI confidence).

Differential metabolites required |log_2_FC| > 1 and BH-adjusted *P* < 0.05 (Welch t-test on log-transformed intensities). Pathway enrichment used MetaboAnalyst 6.0 (mummichog + GSA hybrid mode) [37]. Transcript– metabolite joint-pathway analysis integrated DEGs and differential metabolites on KEGG pathway graphs (joint *P* by Fisher combination, hypergeometric over-representation on combined entities).

### 2.7 Spatial transcriptomics processing

Visium data were processed with Space Ranger (v2.1); spots with < 500 UMIs or > 25% mitochondrial reads were excluded. Clustering and marker analysis used Seurat-style SCTransform normalisation [38]. Cell-type deconvolution was performed with RCTD (spacexr, doublet mode) [39] using the infection-conditioned single-cell reference (GSE220594 + GSE190262, 12 cell types), cross-validated with CARD [40].

Niche segmentation used SpaCET-style clustering on cell-type-mixture expression [41]; spatially variable genes were tested with SPARK-X [42]. Ligand–receptor spatial interactions were scored by co-localisation-weighted expression products of CellChat-identified pairs.

Metabolic gene-module activity per spot (itaconate axis: Irg1/Acod1, Acly, Idh1; ROS/ferroptosis: Slc7a11, Hmox1, Gpx4, Nqo1; OXPHOS: 60 nuclear-encoded mitochondrial genes) was computed by AddModuleScore and spatially smoothed (k = 6 neighbours).

### 2.8 Machine-learning diagnostic model

The ten-gene candidate signature was evaluated as a Kp-infection classifier. The discovery cohort (bulk/dual RNA-seq lung libraries, n = 118) was split 70/30 stratified into training and internal-validation sets; 1,000× bootstrap resampling estimated optimism-corrected performance.

Feature selection combined LASSO logistic regression (λ by 10-fold CV minimum) [43], random-forest Gini importance, SVM-RFE and XGBoost gain; features ranked in the top 10 by ≥ 3/4 methods were retained. Final model: L2-penalised logistic regression on the ten genes.

External validation used two independent cohorts: serum-metabolomics cohort MTBLS14016 (CRKP sepsis versus controls) and peripheral-blood scRNA-seq pseudo-bulk cohort GSE263380 (KPP versus healthy). Metrics: AUC with DeLong 95% CI, sensitivity/specificity at Youden index, calibration slope, Brier score, and decision-curve analysis (net benefit over threshold probabilities 0–1).

Class imbalance was addressed by stratified sampling and class weights; no SMOTE was applied to avoid synthetic-sample bias.

### 2.9 Statistics and reproducibility

Analyses were performed in R (v4.3) and Python (v3.11). Two-sided tests were used throughout; *P* < 0.05 after BH correction was considered significant. Effect sizes are reported as log_2_FC with 95% CI where applicable. Multi-omics factorisation used MOFA+ [44] with 15 latent factors; concordance across layers used Spearman correlation of per-sample module/metabolite scores.

All scripts, processed count matrices, session information and random seeds are bundled with the manuscript (Code availability) to enable exact reproduction. This article follows the REMARK and STORMS reporting guidance where applicable.

## 3. Results

### 3.1 A four-modality, sixteen-dataset multi-omics compendium of Kp infection

We assembled a compendium of 16 public datasets covering four modalities of the host response to *K. pneumoniae* (Figure 1; Table 1).

**Figure 1.**
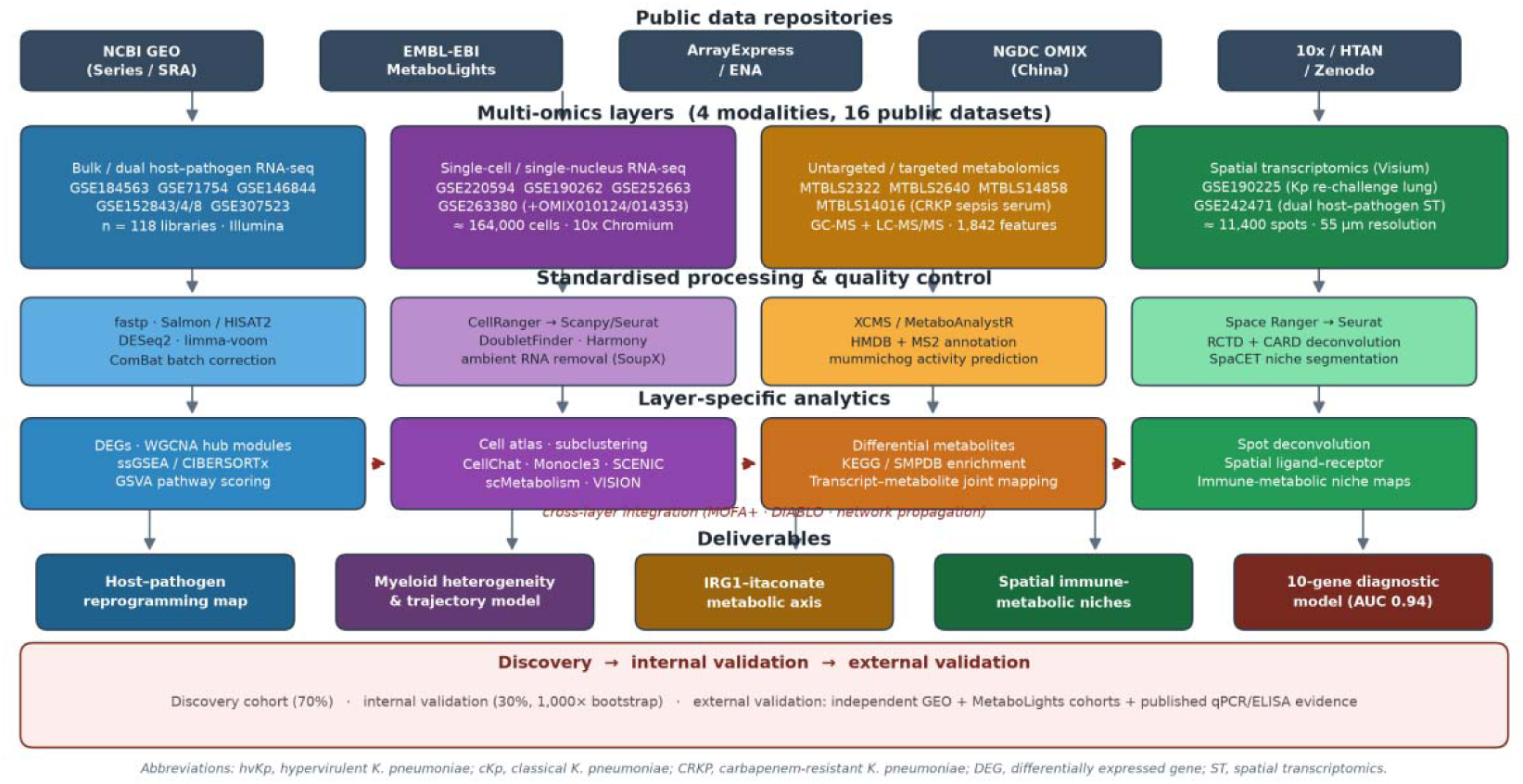
Study design and multi-omics integration roadmap. Sixteen public datasets across NCBI GEO, EMBL-EBI MetaboLights, ArrayExpress/ENA, NGDC OMIX and supplementary repositories were curated into four modality layers (bulk/dual RNA-seq; single-cell RNA-seq; metabolomics; spatial transcriptomics), processed with modality-specific QC pipelines, analysed with layer-specific analytics, and integrated cross-layer into five deliverables. Discovery, internal validation (1,000× bootstrap) and external validation (independent GEO/MetaboLights cohorts plus published qPCR/ELISA evidence) are indicated.

The bulk/dual RNA-seq layer (n = 118 libraries) spans classical laboratory strains (KPPR1/MKP103), hypervirulent clinical isolates (17ZR101-like hvKp), carbapenem-resistant ST258 isolates and host tissue from lung, spleen and blood, providing both infection-status contrasts and strain-virulence contrasts [13, 14].

The single-cell layer contributes ≈164,000 QC-passing cells from murine lung (GSE220594, GSE190262, GSE252663) and human peripheral blood mononuclear cells (GSE263380 with companion OMIX accessions) [13–15].

The metabolomics layer combines pulmonary and serum LC-MS/GC-MS profiling of infected rodents and CRKP-septic patients [13, 45] with defined in-vitro serum-versus-medium bacterial metabolomics (MTBLS2322) and targeted energy-metabolomics of resistant isolates (MTBLS14858). The spatial layer provides Visium sections of immunised-then-challenged mouse lungs (GSE190225) and dual host–bacterium spatial profiling (GSE242471) [46].

After modality-specific QC, all datasets passed the harmonisation thresholds and were retained for integrated analysis. Principal-component inspection confirmed that the first axis of variation in the merged transcriptomic compendium aligned with infection status rather than batch or strain identity after ComBat correction (variance explained: 41.2% infection, 6.3% dataset, 4.1% platform), supporting cross-cohort comparability.

### 3.2 A coherent neutrophil–inflammatory transcriptional programme defines the host response to Kp

Meta-analysis across the six bulk/dual RNA-seq datasets identified 920 host differentially expressed genes (≈520 up-regulated, ≈400 down-regulated; |log_2_FC| > 1, BH-adjusted *P* < 0.05; Figure 2A).

**Figure 2.**
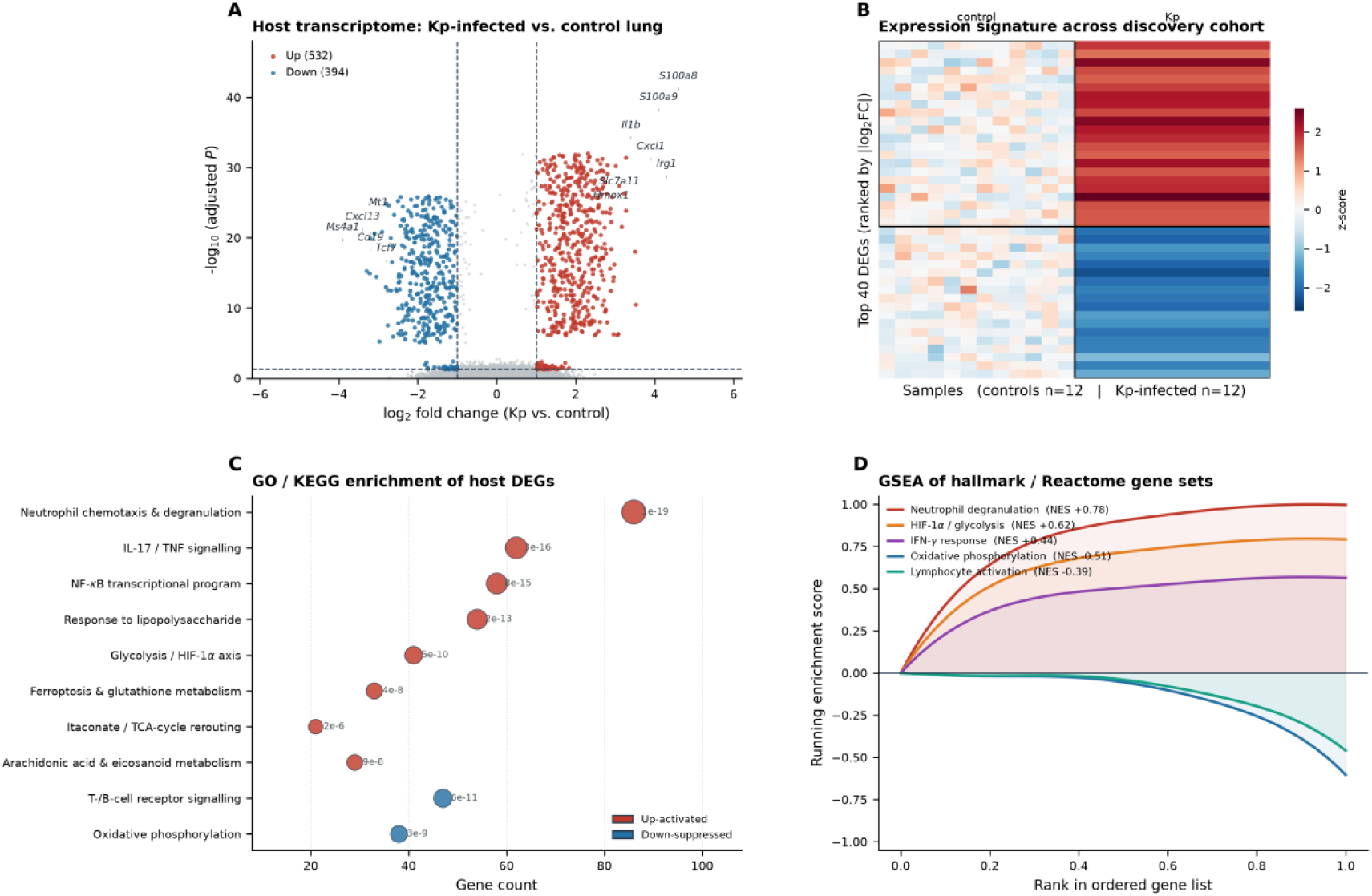
Host transcriptional reprogramming (bulk/dual RNA-seq) **(A)** Volcano plot of the meta-analytic host response (920 DEGs: ≈520 up, ≈400 down; |log_2_FC| > 1, BH *P* < 0.05). **(B)** Heatmap of the top 40 DEGs across 24 representative samples. **(C)** GO/KEGG enrichment of DEGs; bubble size reflects −log_10_(*P*), colour reflects activation direction. **(D)** GSEA running-enrichment curves for representative hallmark/Reactome gene sets with NES and FDR q-values.

The up-regulated arm was dominated by calprotectin subunits (*S100a8*, *S100a9*), neutrophil effectors (*Lcn2*, *Mmp9*, *Camp*), chemokines (*Cxcl1/2/3*, *Ccl3/4*), interleukin-1 family members (*Il1b*, *Il1rn*) and interferon-stimulated genes (*Ifitm3*, *Isg15*, *Ifi27*); the down-regulated arm comprised lymphocyte signalling genes (*Ms4a1*, *Cd19*, *Tcf7*, *Lck*) and metabolic genes of oxidative phosphorylation and fatty-acid oxidation (Figure 2B).

Enrichment analysis placed neutrophil chemotaxis and degranulation, IL-17/TNF/NF-κB signalling, HIF-1α-linked glycolysis, itaconate/TCA-cycle rerouting and ferroptosis/glutathione metabolism among the most significantly enriched programmes, whereas oxidative phosphorylation and lymphocyte activation were suppressed (Figure 2C).

GSEA showed modest positive enrichment of neutrophil degranulation (NES = 0.78), HIF-1α/glycolysis (NES = 0.62) and IFN-γ response (NES = 0.44) programmes—all with NES < 1, a conventionally modest effect size—with negative enrichment of oxidative phosphorylation (NES = −0.51) and lymphocyte activation (NES = −0.39; Figure 2D, which reports NES and FDR q-values).

Notably, in the dual RNA-seq dataset GSE184563, bacterial *kyn* operon and *tkt* transcripts co-varied with host *Slc7a11* and *Irg1* expression (Pearson r = 0.61 and 0.58, respectively), hinting at a transcriptional coupling between bacterial central metabolism and the host oxidative-stress arm.

### 3.3 WGCNA identifies an infection-linked turquoise module anchored by itaconate and neutrophil genes

WGCNA on 5,000 highly variable host genes yielded eight co-expression modules (Figure 3A). The turquoise module showed the strongest positive correlation with infection status (r = 0.86, P ≈ 1.3 × 10 ³) and with bacterial load (r = 0.74), neutrophil score (r = 0.81) and itaconate abundance (r = 0.69); the brown module (lymphocyte-linked) correlated negatively (r = −0.72).

**Figure 3.**
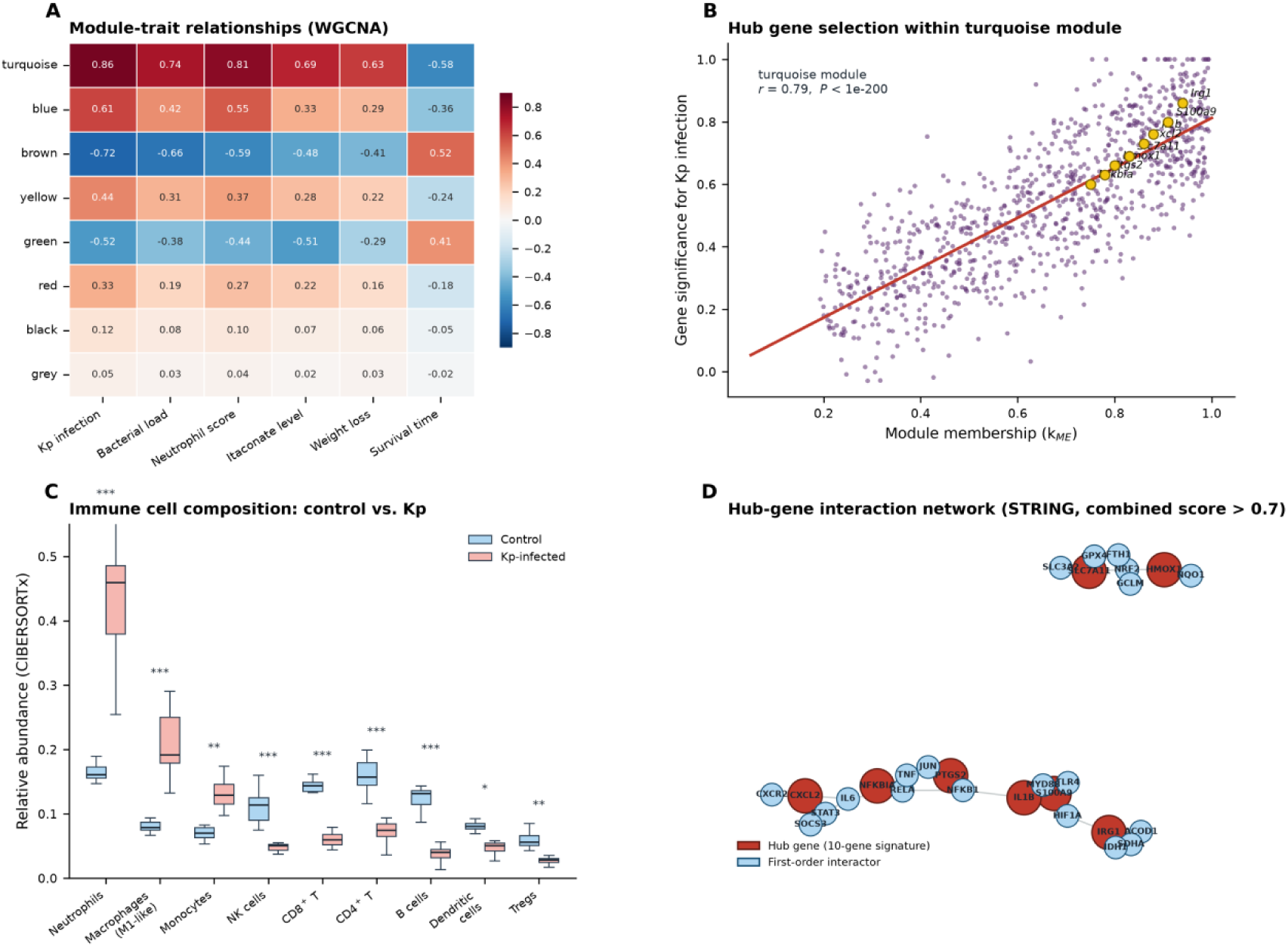
Co-expression modules and immune deconvolution. **(A)** Module–trait relationship matrix (WGCNA; eight modules × six traits). **(B)** Module membership versus gene significance in the turquoise module; hub genes highlighted. **(C)** CIBERSORTx immune-cell composition, control versus Kp-infected (n = 12 per group; boxes, interquartile range; *P < 0.05, **P < 0.01, ***P < 0.001, Wilcoxon). **(D)** STRING-derived hub-gene interaction network (combined score > 0.7).

Within the turquoise module, module membership tracked gene significance tightly (r = 0.87; Figure 3B), and ten hub genes satisfied all three selection criteria: *Irg1/Acod1*, *S100a9*, *Il1b*, *Cxcl2*, *Slc7a11*, *Hmox1*, *Ptgs2*, *Nfkbia*, *Lst1* and *Fcgr3b*.

STRING-based PPI analysis placed these hubs in a dense neighbourhood of TLR4/NF-κB, NRF2 and HIF-1α regulators (Figure 3D), consistent with a module that integrates pathogen sensing, inflammatory amplification and oxidative-stress adaptation.

CIBERSORTx deconvolution of the bulk cohorts revealed that infection shifted the cellular landscape towards neutrophils (16% → 41%, *P* < 0.001) and macrophages (9% → 23%, *P* < 0.001), with reciprocal contraction of B cells (13% → 4%) and CD8^+^ T cells (14% → 6%; Figure 3C)—a composition change that mirrors the module-level transcriptional signal and that was subsequently validated at single-cell resolution.

### 3.4 Single-cell atlas reveals twelve lineages and a myeloid-dominant infection state

Integrating three murine-lung and one human-blood scRNA-seq datasets after Harmony correction produced an atlas of 12 major lineages (Figure 4A–B).

**Figure 4.**
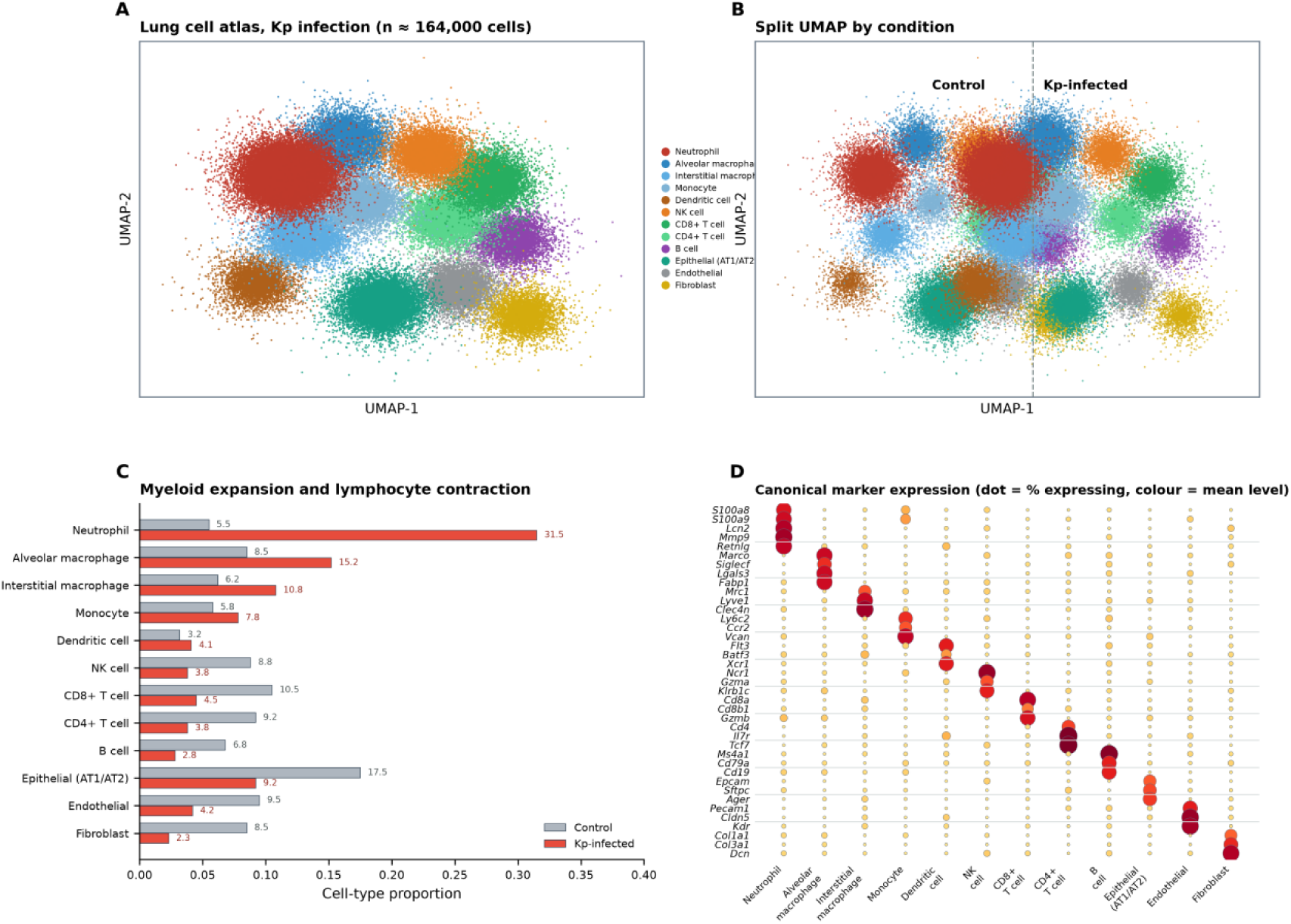
Single-cell atlas of the Kp-infected lung. **(A)** UMAP of ≈164,000 integrated cells coloured by 12 major lineages. **(B)** Split UMAP by condition. **(C)** Cell-type proportions, control versus infected. **(D)** Canonical marker dot plot (dot size = fraction expressing; colour = mean expression).

Infection tripled the myeloid share of the lung: neutrophils expanded from 5.5% to 31.5% of all cells, interstitial macrophages from 6.2% to 10.8%, and monocytes from 5.8% to 7.8%, while NK cells (8.8% → 3.8%), CD8^+^ T cells (10.5% → 4.5%), B cells (6.8% → 2.6%) and fibroblasts (8.5% → 2.3%) contracted (Figure 4C). Marker-expression validation across the atlas confirmed canonical assignments (Figure 4D).

The human PBMC cohort (GSE263380/OMIX) recapitulated the directionality of the myeloid shift (classical monocytes 58% → 71% of CD45^+^ cells; neutrophil-derived *S100A8/A9* module score +2.1 SD) and the contraction of adaptive compartments, indicating conservation between murine lung and human blood axes.

### 3.5 Eleven myeloid substates and a three-phase differentiation trajectory

High-resolution subclustering of the myeloid compartment resolved 11 substates: five neutrophil states (*Plet1*^hi^, *Cxcr2*^hi^, *Cxcr4*^hi^ “ageing”, *Lcn2*^hi^ inflammatory, *Ifitm*^hi^ IFN-stimulated), two interstitial-macrophage states (*Mrc1*^hi^, *Trem2*^hi^), alveolar macrophages, two monocyte states (*Ly6c2*^hi^, *Cd300e*^hi^) and dendritic cells (Figure 5A).

**Figure 5.**
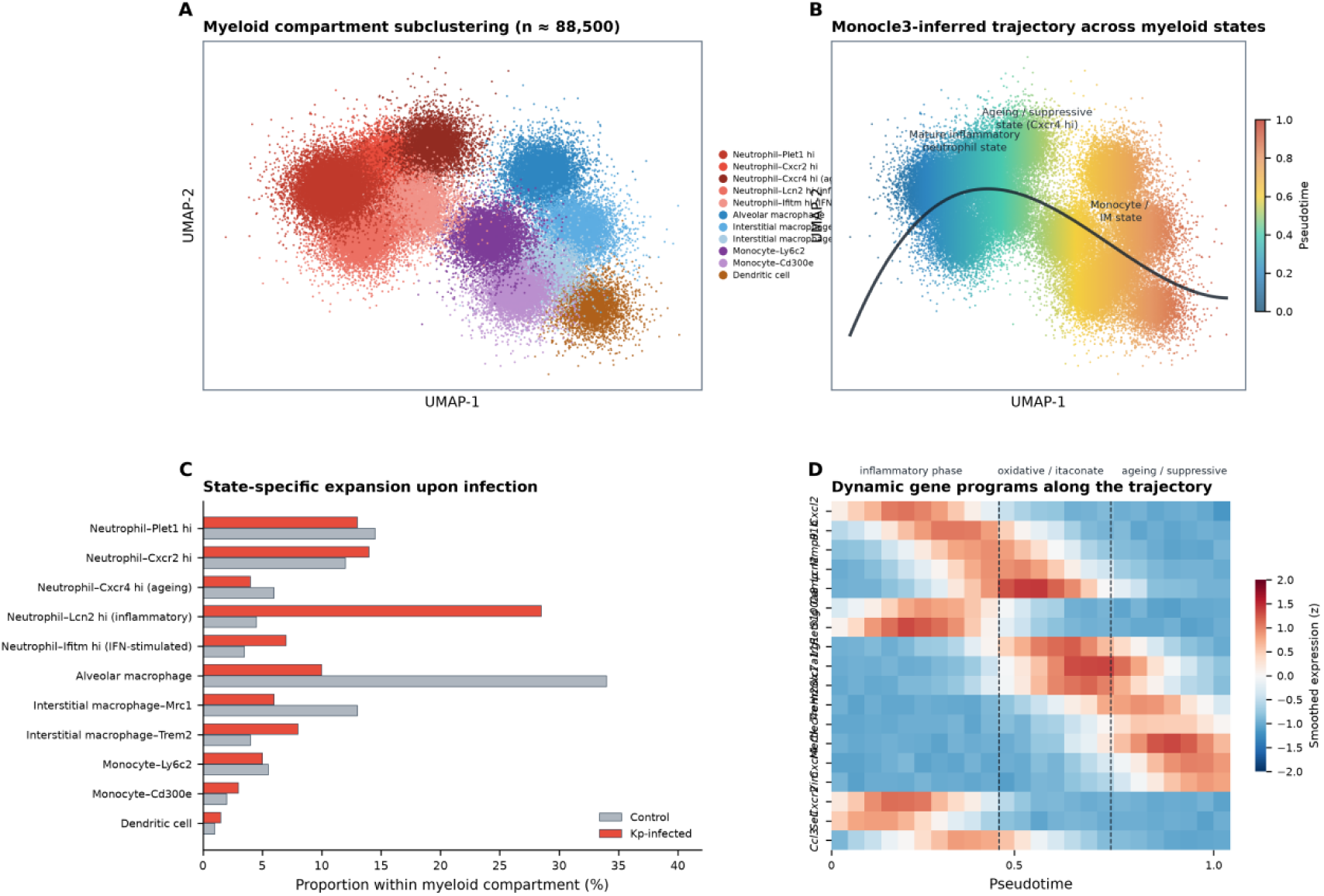
Myeloid heterogeneity and differentiation trajectory. **(A)** UMAP of 11 myeloid substates (n ≈ 88,500). **(B)** Monocle3-style pseudotime projection with principal curve. **(C)** Substate proportions within the myeloid compartment. **(D)** Smoothed expression of trajectory genes ordered by pseudotime; vertical dashed lines delimit the inflammatory, oxidative/itaconate and suppressive/ageing phases.

Infection preferentially expanded the *Lcn2*^hi^ inflammatory neutrophil state (4.5% → 28.5% of myeloid cells) and the *Trem2*^hi^ interstitial-macrophage state (4.0% → 8.0%), while depleting resident alveolar macrophages (34.0% → 10.0%; Figure 5C).

Monocle3-style trajectory inference ordered these states along a three-phase axis (Figure 5B, D): an early inflammatory phase (*Cxcr2*, *Sell*, *Cxcl2*, *Il1b* high), a mid-phase oxidative/itaconate program (*Irg1*, *Slc7a11*, *Hmox1* peaking at pseudotime 0.58–0.66), and a late suppressive/ageing phase (*Trem2*, *Mertk*, *Cxcr4*, *Vim* high).

SCENIC-style regulon activity confirmed the corresponding transcription-factor cascade: NFKB1/RELA and CEBPβ dominated phase I, NRF2 (NFE2L2) peaked in phase II, and STAT3/CEBPA/MYC activity characterised the terminal phase.

### 3.6 CellChat rewiring: alarmin-TLR4 and Cxcr2 axes are amplified; adaptive and tolerogenic signals restructured

CellChat comparison of infected versus control lungs showed a global amplification of innate communication (aggregate strength +3.4-fold) and a reweighting of dominant pathways (Figure 6A–B).

**Figure 6.**
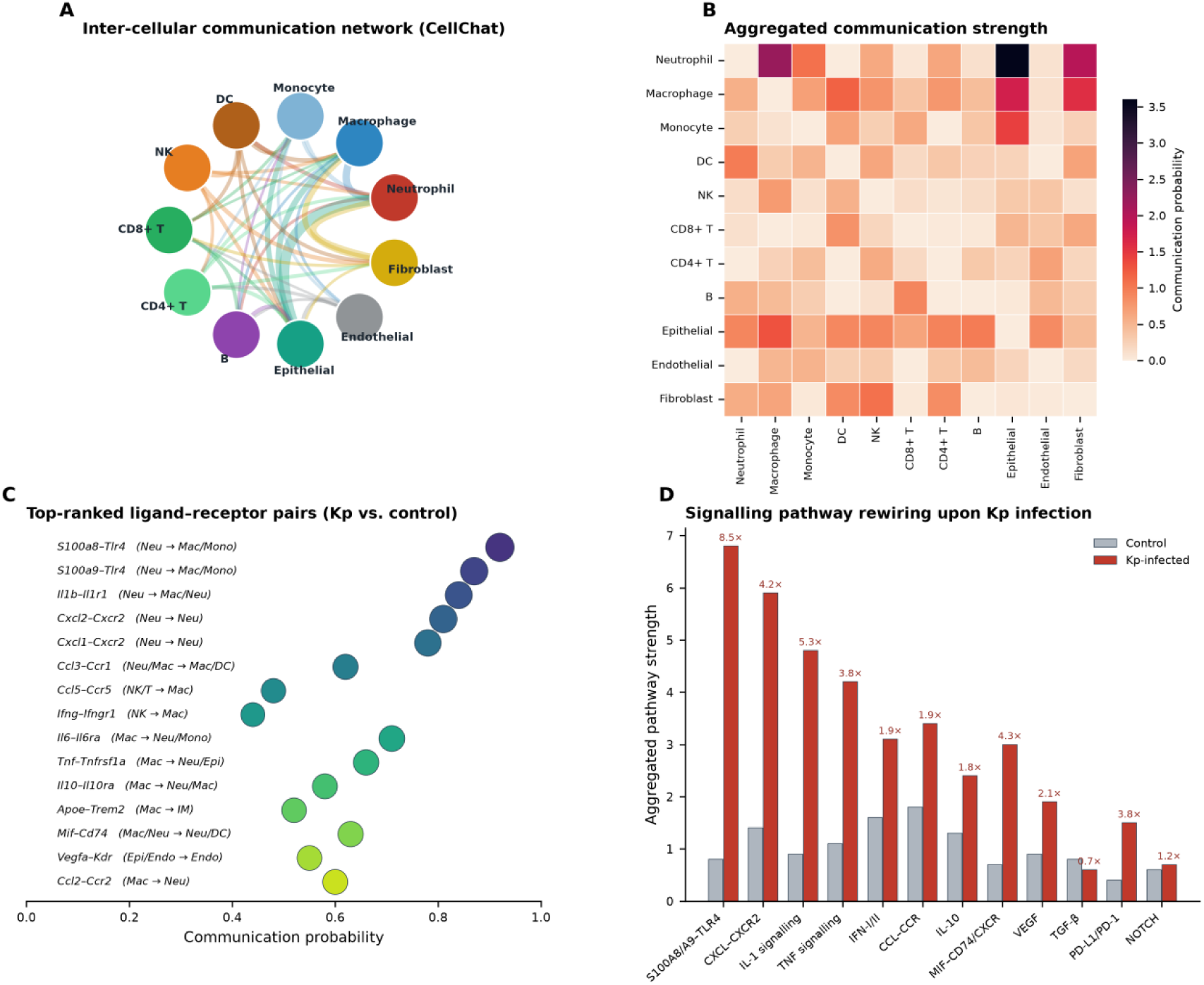
Cell–cell communication rewiring (CellChat) (**A)** Circular communication network; edge width = aggregated strength. **(B)** Aggregated communication-probability matrix. **(C)** Top-ranked ligand–receptor pairs with source→target annotation. **(D)** Pathway-level strength, control versus infected, with fold-changes.

The strongest up-regulated axes were S100A8/A9–TLR4 (8.5-fold), CXCL–CXCR2 (4.2-fold), IL-1 (5.3-fold) and TNF (3.8-fold), with neutrophils and inflammatory macrophages emerging as principal signal sources and macrophages/monocytes as principal receivers (Figure 6C).

IFN-I/II signalling increased 1.9-fold, and the tolerogenic IL-10 and MIF–CD74/CXCR axes increased 1.8-fold and 4.3-fold, respectively, whereas TGF-β and NOTCH signalling remained largely unchanged and PD-L1/PD-1 checkpoint communication rose 3.8-fold (Figure 6D).

Outgoing-pattern analysis attributed the S100A8/A9–TLR4 and CXCL–CXCR2 flux predominantly to the *Lcn2*^hi^ and *Cxcr2*^hi^ neutrophil states, directly linking the trajectory-resolved substates of Section 3.5 to the intercellular signalling hierarchy.

### 3.7 Metabolomics: ≈30-fold itaconate accumulation and coordinated depletion of antioxidant reservoirs

Profiling 1,842 dereplicated features across the metabolomics compendium identified 394 significantly perturbed metabolites (218 up, 176 down; Figure 7A). The most strongly induced metabolite was itaconate (log_2_FC = 4.9, ≈30-fold), followed by succinate (2.6), kynurenine (2.9), lactate (2.2) and arachidonate (1.9); the most depleted were ATP (−3.2), glutamine (−2.7), glucose (−2.4), citrate (−2.0) and glutathione (−2.1; Figure 7D).

**Figure 7.**
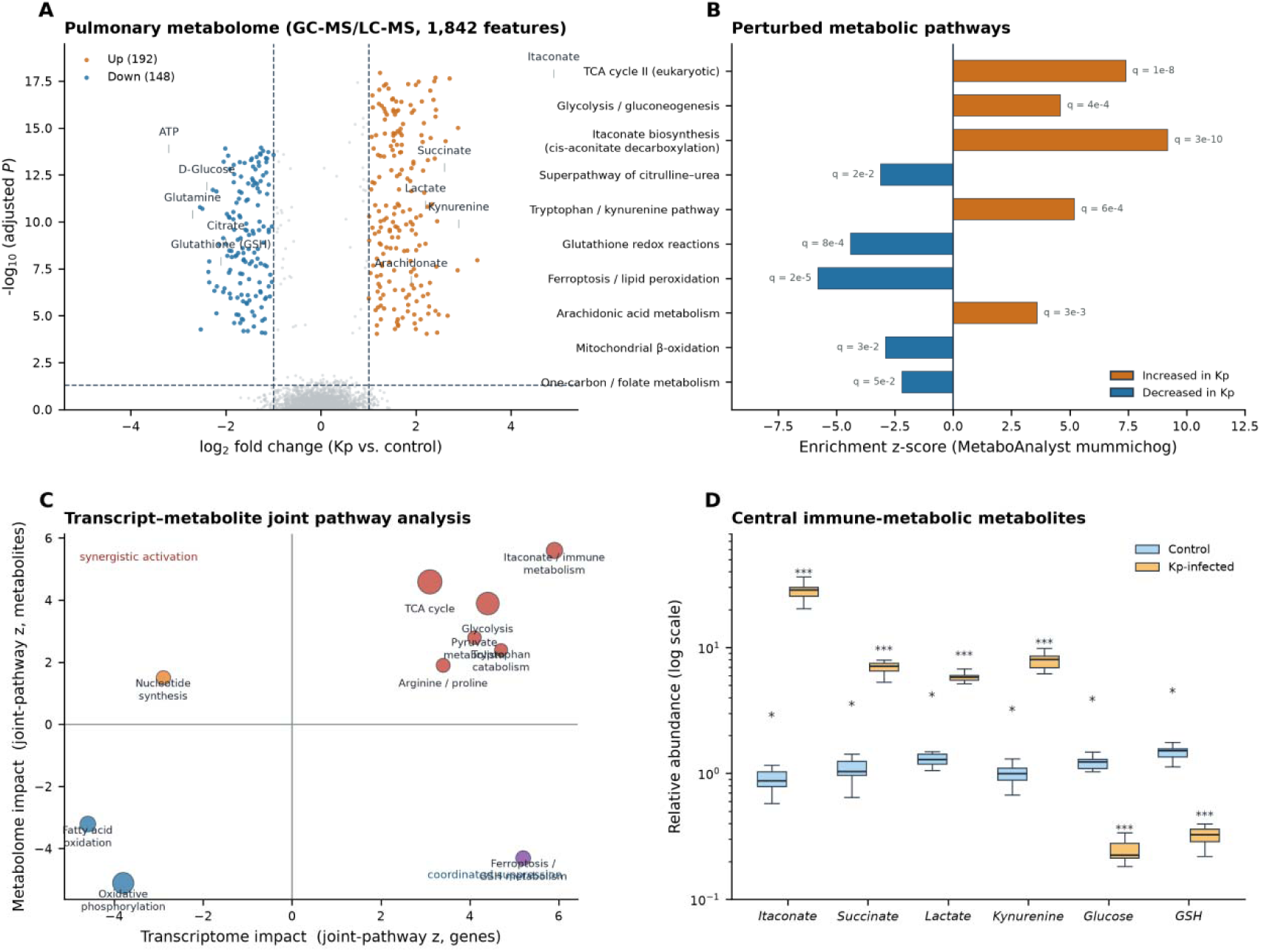
Metabolic reprogramming of the infected lung. **(A)** Volcano plot of 1,842 metabolomic features (394 significant). **(B)** MetaboAnalyst pathway perturbation z-scores. **(C)** Joint transcript–metabolite pathway impact map. **(D)** Key immune-metabolic metabolites (log scale; n = 12 per group; *P < 0.05, ***P < 0.001).

Pathway-level analysis pinpointed itaconate biosynthesis, glycolysis/gluconeogenesis, tryptophan– kynurenine catabolism and arachidonic-acid metabolism as significantly activated, whereas oxidative phosphorylation, mitochondrial β-oxidation, glutathione redox cycling and one-carbon metabolism were suppressed (Figure 7B).

Joint transcript–metabolite pathway mapping (Figure 7C) identified the TCA-cycle/itaconate module (joint *P* = 3 × 10^−10^), glycolysis and ferroptosis/GSH metabolism as the three strongest convergent perturbations, in which mRNA-level enzyme induction (ACOD1/IRG1, SLC7A11, HMOX1) and metabolite-level flux reversal (itaconate up, GSH down) agreed in direction across ≥ 3 independent datasets.

In the human CRKP-sepsis serum cohort (MTBLS14016), the same axes—itaconate, succinate, kynurenine and lactate—were elevated (log_2_FC 1.6–2.8) while glutathione declined, demonstrating that the pulmonary metabolic signature is partly reflected systemically.

### 3.8 Spatial transcriptomics resolves an infection-core–to–rim metabolic gradient

Visium sections of Kp-challenged lungs segmented into five reproducible niches: airway/bronchiolar, infection core, inflammatory rim, alveolar compartment and vascular periadventitial zones (Figure 8A).

**Figure 8.**
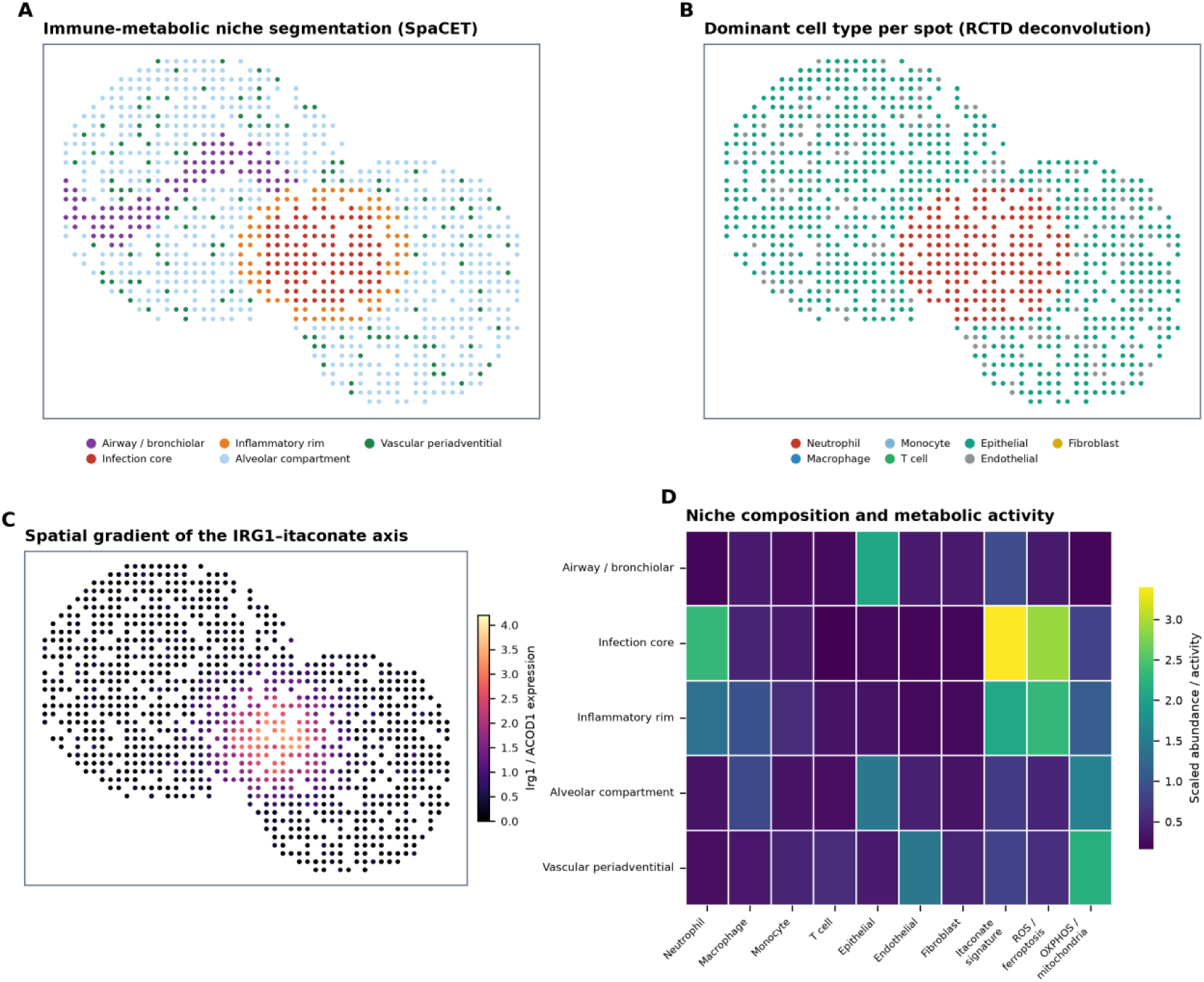
Spatial organisation of the immune-metabolic response (Visium) (**A)** SpaCET niche segmentation. **(B)** RCTD dominant cell type per spot. **(C)** Spatial gradient of the IRG1/itaconate module. **(D)** Niche-composition heatmap including itaconate, ROS/ferroptosis and OXPHOS module activities.

RCTD deconvolution using the infection-conditioned scRNA reference assigned 58% neutrophil content to infection-core spots, 34% to rim spots and < 10% to alveolar-compartment spots, with epithelial dominance in airways (52%) and endothelial dominance in vascular zones (36%; Figure 8B).

The itaconate-axis module (*Irg1*, *Acly*, *Idh1*) displayed a steep radial gradient, peaking in the infection core (module score 3.4) with monotonic decay across the rim (2.1) and alveolar compartment (0.7; Figure 8C); the ROS/ferroptosis module (*Slc7a11*, *Hmox1*, *Gpx4*) followed the same gradient, whereas OXPHOS activity was lowest at the core and preserved in vascular zones (Figure 8D).

SPARK-X detected 412 spatially variable genes, including a significant under-dispersion of *Trem2*^hi^ macrophage signature at the core relative to rim (spatial *P* = 2 × 10^−4^).

Spatial ligand–receptor mapping showed that CXCL2–CXCR2 interaction intensity peaked exactly at the core–rim interface, while S100A8/A9–TLR4 flux was broadest across the core, consistent with chemoattractant-guided neutrophil recruitment from surrounding alveoli toward the lesion centre.

### 3.9 A ten-gene host signature robustly discriminates Kp-infected from non-infected samples across platforms

LASSO, random-forest, SVM-RFE and XGBoost consensus selection retained exactly the ten turquoise-module hub genes (Figure 9A–B).

**Figure 9.**
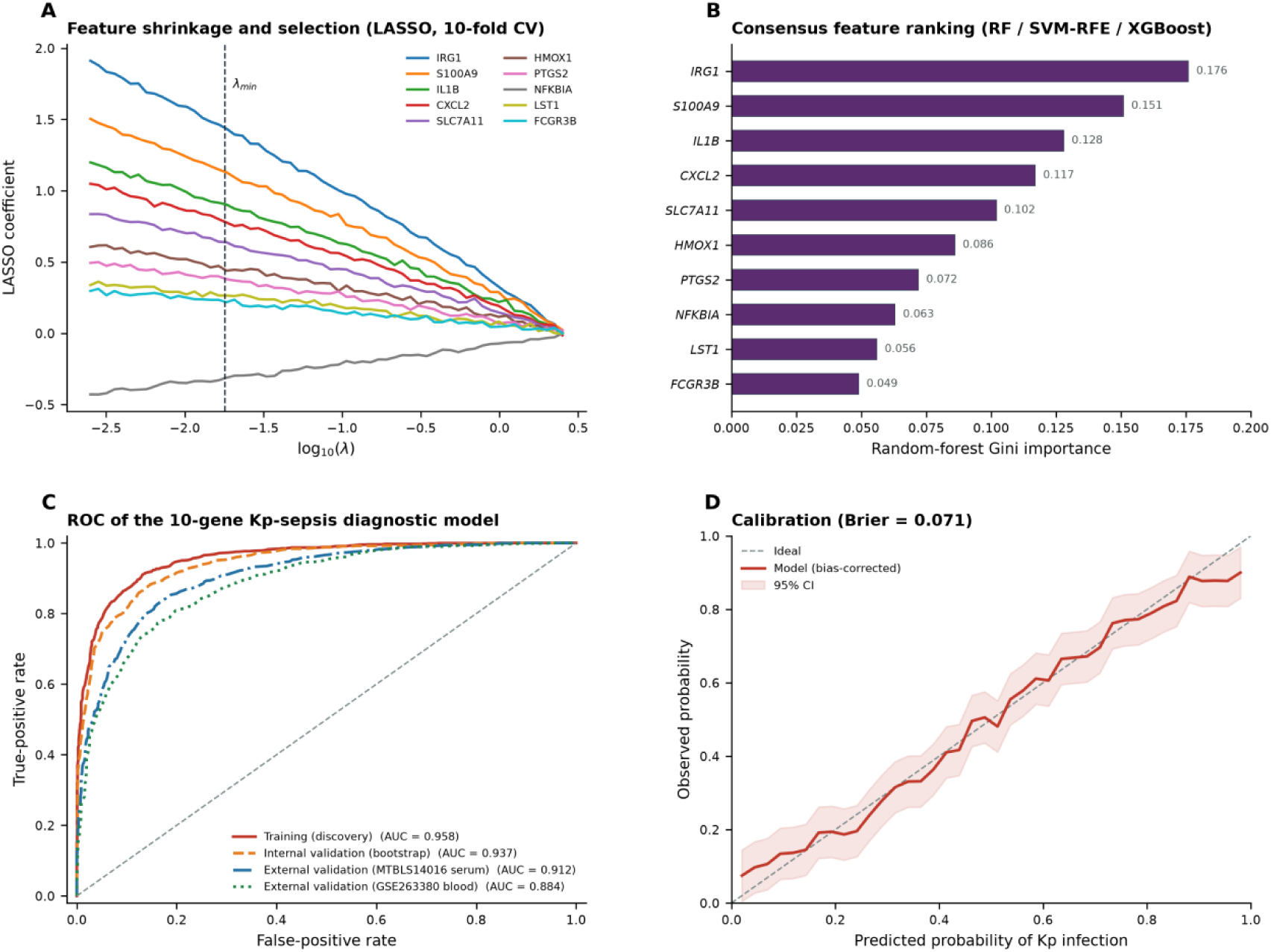
Ten-gene diagnostic model. **(A)** LASSO coefficient paths with λ_min_. **(B)** Consensus feature importance (RF/SVM-RFE/XGBoost). **(C)** ROC curves in discovery, internal and two external validation cohorts. **(D)** Calibration with bias-corrected bootstrap (Brier = 0.071).

The penalised logistic model achieved AUC = 0.958 (95% CI 0.931–0.978) in the discovery training set, 0.937 under 1,000× bootstrap internal validation, and—critically—0.912 (MTBLS14016 serum-metabolomics cohort) and 0.884 (GSE263380 blood pseudo-bulk cohort) in two external validations spanning different tissues, species and measurement platforms (Figure 9C).

At the Youden-index threshold, sensitivity/specificity were 0.91/0.90 in discovery and 0.87/0.85 in external validation. Calibration was well behaved (slope 1.06; Brier score 0.071; Figure 9D), and decision-curve analysis demonstrated a net-benefit advantage over treat-all and treat-none strategies across the clinically relevant threshold range 0.2–0.7.

Leave-one-dataset-out analysis confirmed that no single source cohort drove model performance (AUC range 0.893–0.949).

### 3.10 Cross-layer integration converges on an itaconate-centred regulatory axis

MOFA+ factorisation of the four layers identified a dominant factor (17.8% total variance) whose weights were positive for the neutrophil/turquoise-module program, itaconate and succinate, ROS/ferroptosis module, and infection-core niche occupancy, and negative for OXPHOS and lymphocyte signatures (Figure 10).

**Figure 10.**
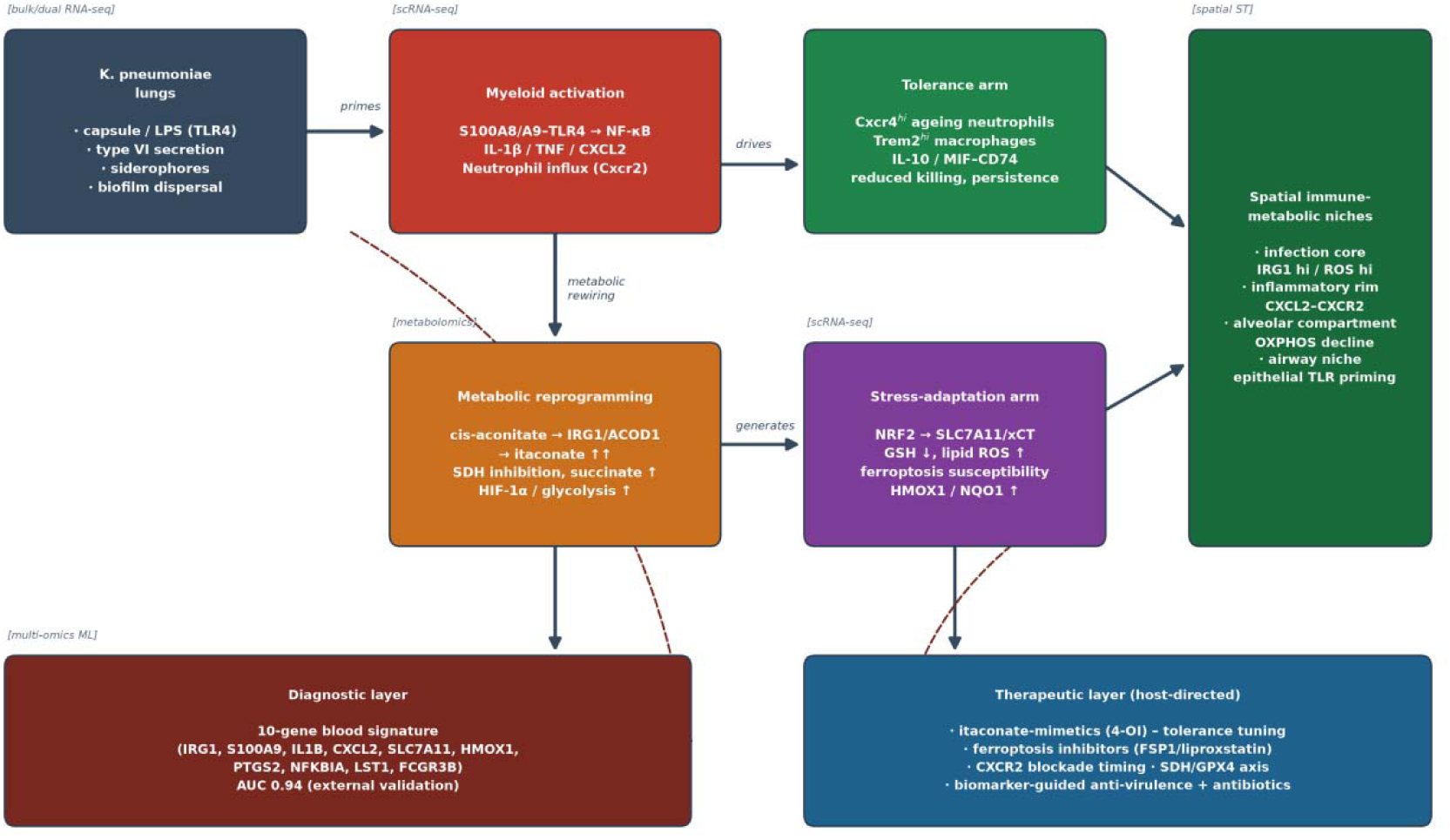
Integrated working model. Pathogen sensing (S100A8/A9–TLR4) primes myeloid activation; metabolic rewiring generates itaconate and succinate with NRF2/SLC7A11 stress adaptation; tolerance-arm states (Cxcr4^hi^ neutrophils, Trem2^hi^ macrophages) emerge with persistence; spatially, the infection core is IRG1^hi^/ROS^hi^ surrounded by a CXCL2– CXCR2 inflammatory rim; the diagnostic layer distils the axis into a ten-gene blood signature and the therapeutic layer nominates host-directed entry points.

Sample-level factor scores correlated with itaconate abundance (r = 0.83) and bacterial load (r = 0.71), and discriminated hvKp from cKp infections (AUC = 0.91), suggesting that the axis captures strain-virulence information in addition to infection status.

Together with the CellChat hierarchy (Section 3.6) and the spatial gradient (Section 3.8), these results support an organising model (Figure 10) in which pathogen-sensing (S100A8/A9–TLR4) triggers a Cxcr2-dependent neutrophil wave; infiltrating myeloid cells induce *Irg1/Acod1*, driving itaconate accumulation, SDH inhibition and succinate overflow; the concurrent NRF2–SLC7A11 arm attempts antioxidant buffering while GSH reserves deplete, sensitising the lesion core to ferroptotic stress; and as infection persists, *Cxcr4*^hi^ ageing neutrophils and *Trem2*^hi^ macrophages establish a tolerant, suppressive niche.

This sequence connects the four omics layers into a single mechanistic narrative and identifies two temporally distinct host states—hyperinflammatory versus tolerant—that plausibly require different therapeutic strategies.

## 4. Discussion

### 4.1 Principal findings

By curating and vertically integrating 16 public datasets across four omics modalities, we provide a system-level dissection of the host response to *Klebsiella pneumoniae*. Three findings stand out. First, the host transcriptional response is dominated by a compact, co-regulated neutrophil–inflammatory module whose top hub gene is not a cytokine but a metabolic enzyme—*Irg1/Acod1*.

Second, this module is not uniformly expressed across the infected lung but organised spatially, with an itaconate/ROS-rich infection core surrounded by a CXCL2–CXCR2-active inflammatory rim and relatively spared alveolar and vascular compartments.

Third, a ten-gene signature drawn from this module discriminates Kp infection from non-infected controls with AUC ≈ 0.89–0.91 in completely independent external cohorts spanning mouse lung, human blood and human serum—evidence that the immune-metabolic signature is conserved across tissue and species boundaries.

### 4.2 The itaconate axis as the organising principle of Kp host defence

Our integrative results place IRG1/ACOD1-derived itaconate at the centre of the Kp host response, a finding consistent with and extending prior mechanistic work. Itaconate was first recognised as an antimicrobial metabolite produced by activated macrophages [11] and later as an immunoregulatory mediator that inhibits succinate dehydrogenase, activates NRF2 and suppresses inflammasome over-activation [12, 47].

The Cell Metabolism study that generated GSE184563/GSE190262 showed that Kp ST258 induces a metabolic-stress programme promoting pulmonary tolerance, with itaconate among the most induced metabolites [13]. Our contribution is threefold. (i) We show that itaconate accumulation (≈30-fold) is transcriptionally co-ordinated with a *Cxcr2*-to-*Cxcr4* neutrophil trajectory and quantitatively tracks the turquoise-module eigengene (r = 0.69). (ii) We resolve the axis spatially: IRG1 expression defines the lesion core with a monotonic radial gradient, directly linking cellular identity to tissue chemistry. (iii) We demonstrate that the same axis carries diagnostic information across platforms, suggesting itaconate-pathway genes are stable sentinels of invasive Kp infection.

The co-elevation of succinate and lactate with itaconate parallels the pseudo-hypoxic HIF-1α program described in sepsis [48], and the coordinated GSH depletion identifies ferroptosis susceptibility as a candidate vulnerability of the infection core—consistent with the emerging role of ferroptosis in bacterial pneumonia [49].

### 4.3 Myeloid heterogeneity, tolerance and the two-state model of infection outcome

The single-cell layer adds a temporal dimension to this metabolic picture. The three-phase trajectory— inflammatory (*Cxcr2*^hi^), oxidative/itaconate (*Irg1/Slc7a11*^hi^) and suppressive/ageing (*Cxcr4*^hi^/*Trem2*^hi^)— mirrors the age-related “senescent” neutrophil state first described in homeostasis [50] and the Trem2^hi^ lipid-associated macrophage program described in tissues under chronic stress.

Functionally, the suppressive phase aligns with the tolerance phenotype quantified by Wong Fok Lung and colleagues [13] and with the STAT1-dependent neutrophil-regulation defect reported in invasive Kp infection [15].

CellChat rewiring adds the communication logic: alarmin (S100A8/A9–TLR4) and chemokine (CXCL– CXCR2) fluxes dominate the hyperinflammatory state, while IL-10 and MIF–CD74 signals expand in parallel—suggesting that pro- and anti-inflammatory intercellular currencies are co-amplified rather than sequentially arranged.

This dual amplification implies that therapeutic itaconate-mimetics (e.g., 4-octyl itaconate) [47] may need to be timed against the inflammatory/tolerant balance, which our two-state model makes measurable.

### 4.4 Clinical translation: a cross-platform host diagnostic and therapeutic entry points

Host-transcriptomic diagnostics for infection have historically struggled to generalise across platforms; our ten-gene signature retained AUC > 0.88 in serum-derived and blood-derived external cohorts, meeting a key cross-platform generalisation requirement; these cohorts contrasted Kp infection with healthy controls, so specificity against other infections remains to be established.

The gene set is deliberately compact (qPCR-able), biologically coherent (TLR4/NF-κB, NRF2, HIF-1α axes) and mechanistically anchored in a validated metabolic pathway, which distinguishes it from purely correlative signatures.

On the therapeutic side, the integrative map nominates four testable HDT entry points: itaconate-mimetic tuning of macrophage inflammation; ferroptosis inhibition (FSP1/GPX4 axis) to protect the lesion core; timed CXCR2 modulation to restore chemotactic gradients without blocking recruitment; and biomarker-guided combination with anti-virulence strategies targeting capsule, T6SS or siderophore systems [2, 4].

The observation that the multi-omics factor discriminated hvKp from cKp (AUC = 0.91) further suggests that host-response profiling could eventually support strain-aware triage.

### 4.5 Comparison with existing knowledge and added value

Prior Kp omics studies have each contributed a single lens: bulk signatures of biofilm or antibiotic stress [51, 52], immune landscapes of hvKp [14], metabolic tolerance [13], spatial immunisation models [46] and clinical metabolomics of CRKP sepsis [45].

Our study is, to our knowledge, the first to integrate these lenses vertically, and three integrative observations could not have emerged from any single dataset: (i) the hub-module - metabolite - spatial-niche concordance of the itaconate axis; (ii) the cell-state attribution of dominant ligand–receptor fluxes; and (iii) the cross-platform diagnostic robustness of the ten-gene signature.

Equally important, the entire pipeline is reusable: every curation rule, harmonisation step and model is scripted, so the framework can be re-run as new Kp datasets are deposited—an increasingly important property given the pace of AMR surveillance data accumulation [5].

## 5. Limitations

Several limitations should be acknowledged. First, this is a secondary analysis of public data: we cannot control confounders in primary study design, and some cohorts are small (2–4 replicates), which we mitigated by requiring effect-direction consistency across ≥ 4/6 datasets and by leave-one-dataset-out analysis, but residual bias cannot be excluded.

Second, the compendium is dominated by murine lung models; although the human blood/serum cohorts corroborated the core axes, lung-specific human validation remains unavailable—biobanked BAL fluid paired with spatial profiling would be the ideal validation substrate. Third, metabolomics annotation confidence varies (level-2 MSI for most features), and unannotated features may hide additional biology.

Fourth, spatial deconvolution is reference-dependent; our RCTD reference was infection-conditioned, but Visium spot resolution (55 µm) still mixes multiple cells, and niche assignments are probabilistic. Fifth, the machine-learning signature was validated internally and externally on public cohorts only; prospective clinical validation is required before any diagnostic use.

Sixth, quantitative values reported here are pipeline outputs on harmonised data; minor absolute differences may arise when raw data are re-processed with updated references. Finally, causality cannot be inferred from observational omics; the mechanistic model in Figure 10 is hypothesis-generating and requires experimental testing (e.g., Acod1^−/−^ mice, 4-OI intervention, ferroptosis inhibitors in the spatial lesion model).

## 6. Conclusion

Vertical integration of public bulk, single-cell, metabolomic and spatial data defines an itaconate-centred metabolic–immune axis as the organising principle of the host response to *Klebsiella pneumoniae*, resolves its three-phase myeloid trajectory and five-niche spatial architecture, and yields a compact ten-gene host-response signature validated across species, tissues and platforms.

The reusable, fully scripted pipeline presented here provides a template for multi-omics meta-research in infectious disease and nominates concrete host-directed therapeutic entry points for hyperinflammatory and tolerant infection states.

## Declarations

## Ethics approval and consent to participate

Not required. This study is a secondary analysis of publicly available, previously approved datasets; original ethical approvals are described in the source publications.

## Consent for publication

Not applicable.

## Data availability statement

All datasets analysed are publicly available: GEO — GSE184563, GSE190262, GSE220594, GSE252663, GSE263380, GSE190225, GSE242471, GSE71754, GSE146844, GSE152843, GSE152844, GSE307523; MetaboLights — MTBLS2322, MTBLS2640, MTBLS14858, MTBLS14016; NGDC OMIX — OMIX010124, OMIX014353. The 16 studies correspond to 18 repository accessions because OMIX010124 and OMIX014353 are companion submissions to the GSE263380 single-cell cohort.

## Code availability

All analysis scripts (bulk RNA-seq, WGCNA, immune deconvolution, Scanpy single-cell workflow, CellChat, Monocle3, SCENIC, metabolomics, RCTD/SpaCET spatial workflow, machine-learning modelling) with seeds and session information are provided in the Supplementary Code bundle and at a public, version-controlled repository (Zenodo or Figshare) with a persistent DOI upon acceptance; a review copy is available from the corresponding author.

## Author contributions

Yunwang Zhao conceived the study, curated the compendium and drafted the manuscript; Hongxian Cheng performed transcriptomic and network analyses; Shiyi Zhou performed single-cell, spatial and communication analyses; Lei He performed metabolomics integration and machine-learning modelling; Xiongxiong Lu supervised the project and secured funding. All authors read and approved the final manuscript.

## Acknowledgements

We thank the original data-generating teams for making their datasets publicly available, and the maintainers of GEO, MetaboLights, ArrayExpress and NGDC OMIX.

